# Central role of Prominin-1 in lipid rafts during liver regeneration

**DOI:** 10.1101/2022.01.20.477032

**Authors:** Myeong-Suk Bahn, Dong-Min Yu, Myoungwoo Lee, Sung-Je Jo, Ji-Won Lee, Hyun Lee, Arum Kim, Jeong-Ho Hong, Jun Seok Kim, Seung-Hoi Koo, Jae-Seon Lee, Young-Gyu Ko

## Abstract

Prominin-1 (PROM1), a lipid raft protein, is required for maintaining cancer stem cell properties in hepatocarcinoma cell lines, but its physiological roles in the liver have not been well studied. Here, we investigated the role of PROM1 in lipid rafts with a precise molecular mechanism during liver regeneration. We found that the expression of PROM1 increased during liver regeneration after 2/3 partial hepatectomy (PHx) or CCl_4_ injection. Interestingly, hepatocyte proliferation and liver regeneration were attenuated in liver-specific *Prom1* knockout (*Prom1^LKO^*) mice compared to wild-type (*Prom1^f/f^*) mice. Detailed mechanistic studies revealed that PROM1 interacted with the interleukin-6 signal transducer glycoprotein 130 (GP130) and confined GP130 to lipid rafts so that STAT3 signaling by IL-6 was effectively activated. Moreover, the overexpression of the glycosylphosphatidylinsositol (GPI)-anchored first extracellular domain of PROM1 (PROM1^GPI-EX1^), which is a domain that binds to GP130, rescued the proliferation of hepatocytes and liver regeneration in *Prom1^LKO^* mice. PROM1 is upregulated in hepatocytes during liver regeneration, and upregulated PROM1 recruits GP130 into lipid rafts and activates the IL6-GP130-STAT3 axis. Thus, we conclude that PROM1 plays an important role in lipid rafts during liver regeneration and might be a promising target for therapeutic applications of liver transplantation.

## Introduction

The liver has a unique regenerative capability to restore the original liver mass after tissue loss induced by 2/3 partial hepatectomy (PHx) or other liver injuries. PHx is a well-characterized experimental model for liver regeneration in rodents. Mice recover most of their liver mass 7 days after PHx [1]. This regenerative capability is important for maintaining liver function after damage by various factors, including alcohol, viruses, and toxins. During liver regeneration, quiescent hepatocytes proliferate by several cytokines and growth factors, such as epidermal growth factor (EGF), hepatocyte growth factor (HGF), fibroblast growth factor (FGF), tumor necrosis factor-α (TNFα), and interleukin-6 (IL-6) [2, 3].

IL-6 is a pleiotropic cytokine in the body. After PHx or other liver injuries, gut-derived factors such as lipopolysaccharides (LPS) activate Kupffer cells and resident liver macrophages to secrete IL-6 [4]. Secreted IL-6 binds to the interleukin-6 receptor (IL-6R) and then forms a signaling complex consisting of IL-6R and interleukin-6 signal transducer glycoprotein 130 (GP130) in hepatocytes [5]. The complex initiates several downstream signaling pathways, including Janus kinases (JAKs), signal transducer and activator of transcription 3 (STAT3), MAP kinases and the PI3 kinase pathway.

*Il-6* knockout impairs hepatocyte proliferation and induces liver necrosis after PHx in mice, preventing liver mass recovery. As a result, *Il-6* knockout significantly increases mortality after surgery. Thus, a single injection of IL-6 rescues this phenotype in *Il-6* knockout mice [6]. In addition, liver-specific *Stat3* knockout impairs the DNA synthetic response in hepatocytes and decreases the expression of G_1_ phase cyclins such as cyclin D1 and cyclin E [7]. Consistent with the important role of the IL-6 signaling pathway during liver regeneration, liver-specific knockout of suppressor of cytokine signaling 3 (SOCS3), a negative regulator of the STAT3 pathway, exhibits prolonged activation of STAT3 and enhances hepatocyte proliferation, resulting in accelerated liver mass replenishment after PHx [8].

Prominin-1 (PROM1), also known as CD133, is a penta-span transmembrane glycoprotein. PROM1 is associated with distinct detergent-resistant lipid rafts [9] and is found in membrane protrusions such as filopodia and microvilli [10]. PROM1 has been studied as one of the most widely used cancer stem cell (CSC) markers in various human tumors, including the liver [11]. In addition to cancer stem cells, PROM1 is also expressed in normal stem cells, including hematopoietic stem cells and various epithelial cells, in the brain, kidney, digestive track, and liver [12–14]. In addition, PROM1 has been known to regulate the glucagon and TGF-β signaling pathways in the liver by interacting with radixin and SMAD7, respectively [15, 16].

Because PROM1, a marker for hepatic progenitor cells, is also upregulated in hepatocytes after liver injury [16], the upregulated PROM1 might regulate various signaling pathways related to hepatocyte proliferation. Here, we observed a significant increase in the expression of PROM1 in hepatocytes during liver regeneration after PHx or CCl_4_ injection. Liver-specific *Prom1* knockout (*Prom1^LKO^*) mice showed impaired liver regeneration because of reduced hepatocyte proliferation. Mechanistically, we found that the increased PROM1 in hepatocytes confined GP130 to lipid rafts and facilitated activation of STAT3. These results demonstrated that PROM1 plays an important role during liver regeneration through the IL6-GP130-STAT3 signaling pathway.

## Results

### PROM1 is upregulated in hepatocytes during liver regeneration

To investigate the expression of PROM1 during liver regeneration, we performed 2/3 partial hepatectomy (PHx) in wild-type mice. We found that the mRNA level of PROM1 increased after PHx by qRT-PCR (Fig. 1A). The mRNA level of PROM1 peaked 48 hours after PHx and then gradually decreased. Consistently, immunoblotting confirmed that the protein level of PROM1 increased 48 hours after PHx (Fig. 1B). Next, we determined which cells expressed PROM1 in the liver by PROM1 double immunofluorescence with hepatocyte nuclear factor 4α (HNF4α as a specific marker of hepatocytes) or cytokeratin-19 (CK19 as a specific marker of ductal cells) (Fig. 1C, D). PROM1 was mainly expressed in ductal cells of sham liver, whereas it was expressed in hepatocytes of PHx liver.

**Figure 1.**
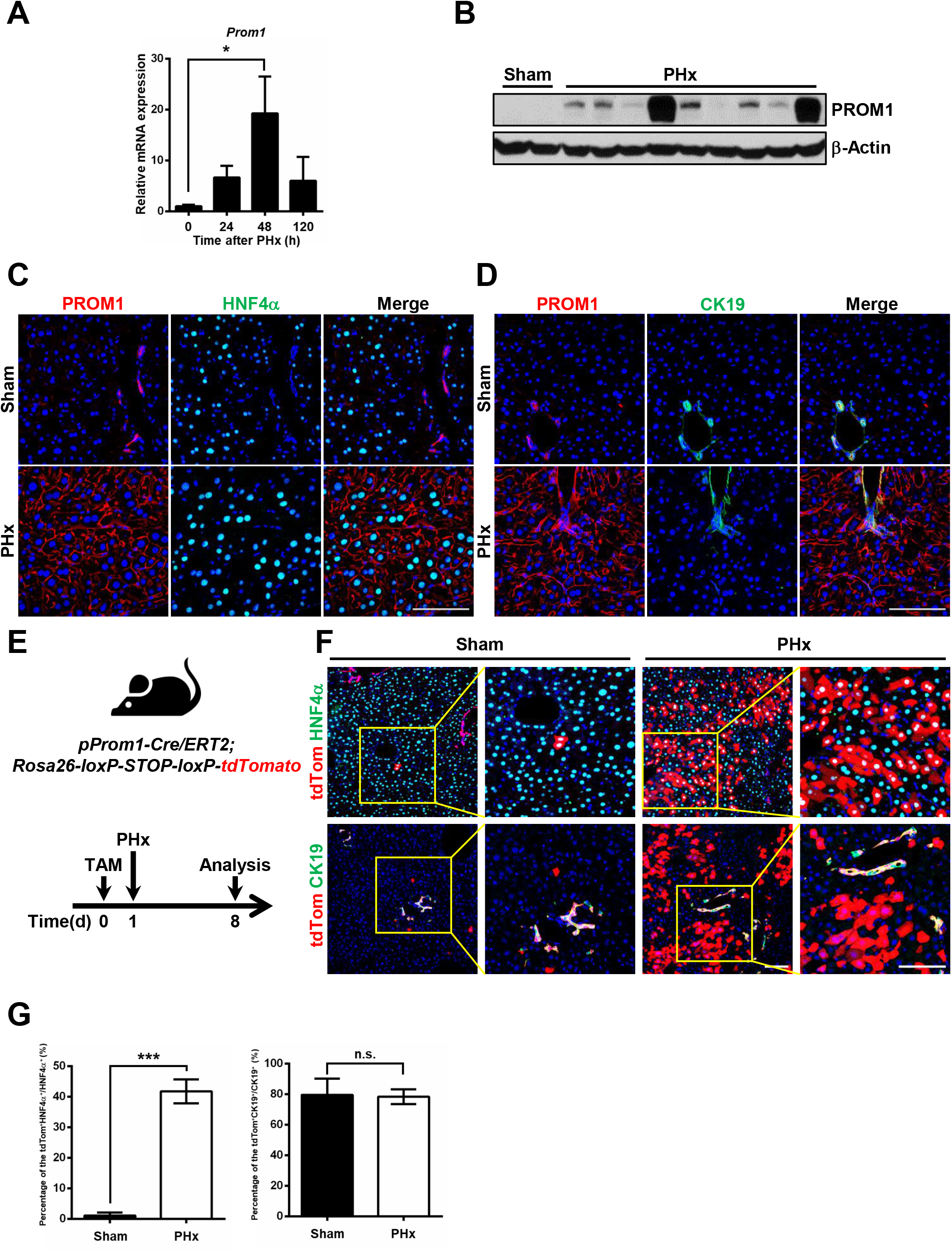
The expression of PROM1 in hepatocytes increases after partial hepatectomy. A 2/3 partial hepatectomy was performed in 8-week-old male wild-type mice. (A) The relative mRNA level of PROM1 in sham and PHx livers (n=3-7). (B) Immunoblotting for PROM1 in wild-type livers 48 hours after PHx. (C, D) Double immunofluorescence for PROM1 and HNF4α (C) or CK19 (D) in sham and PHx livers. (E) *Prom1^Cre/ERT2^; Rosa26^tdTomato^* mice were generated for lineage tracing of cells expressing PROM1 in the liver. PHx was performed 1 day after tamoxifen injection. The mice were analyzed 7 days after sham (n=4) or PHx (n=4). (F) Representative images of tdTom double immunofluorescence with HNF4α or CK19 in sham and PHx livers. (G) The percentage of tdTom-expressing cells was statistically determined from total HNF4α- or CK19-expressing cells. Scale bar = 100 µm. Student *t*-test; \**p* < 0.05, \*\*\**p* < 0.001, n.s., nonsignificant. Data are expressed as the mean ± SEM.

To further clarify the cell types expressing PROM1 during liver regeneration, we generated a lineage tracing mouse in which tdTomato (tdTom) was expressed by tamoxifen in PROM1- positive cells (Fig. 1E) and observed the expression of tdTom after PHx in the liver. Consistent with the immunofluorescence data, the expression of tdTom significantly increased in HNF4α- expressing hepatocytes but not in CK19-expressing ductal cells after PHx (Fig. 1F). Indeed, ∼41% of HNF4a-expressing hepatocytes expressed tdTom (Fig. 1G). These data demonstrate that the expression of PROM1 significantly increases in hepatocytes during liver regeneration after PHx.

### PROM1 deficiency impairs liver regeneration in mice

To determine the role of PROM1 in the process of liver regeneration, we compared the livers of wild-type (*Prom1^f/f^*) and liver-specific *Prom1* knockout mice (*Prom1^LKO^*) after PHx. As a result of measuring the remnant liver-to-body weight ratio following PHx, liver regeneration of *Prom1^LKO^* mice was impaired compared to that of *Prom1^f/f^* mice (Fig. 2A). *Prom1^f/f^* mice recovered their original liver mass almost 5 days after PHx, whereas *Prom1^LKO^* mice did not. Compared with *Prom1^f/f^* mice, the liver-to body weight ratio was significantly lower in *Prom1^LKO^* mice 48 and 120 hours after surgery.

**Figure 2.**
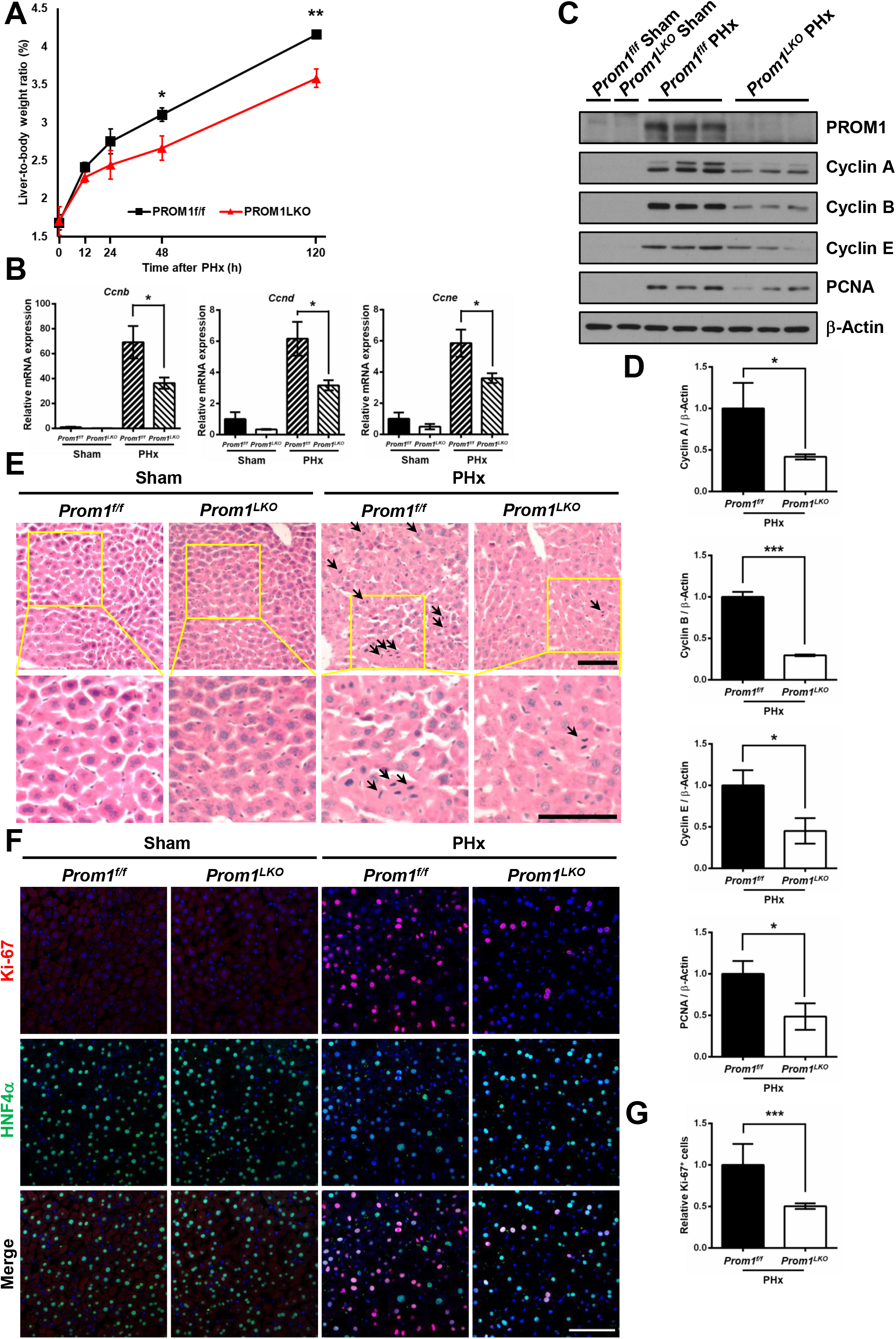
Liver-specific deletion of *Prom1* in mice impairs liver regeneration after partial hepatectomy. A 2/3 partial hepatectomy was performed in 8-week-old male *Prom1^f/f^* and *Prom1^LKO^* mice. (A) Ratio of liver-to-body weight on the indicated days after PHx (n=4-7). (B) The relative mRNA levels of cell cycle genes (*Ccnd, Ccne, Ccnb*) 48 hours after PHx (n=8). Each mRNA level was normalized by 18S rRNA. (C) Immunoblotting for PROM1 and cell cycle proteins (Cyclin A, B, and E, and PCNA) 48 hours after PHx. (D) Statistical analysis of the band intensity in C. The band intensity of each protein was normalized to that of β-actin. (E) Representative H&E staining in the liver 48 hours after PHx. Mitotic cells are indicated by arrows. (F) Representative double immunofluorescence for Ki-67 and HNF4α in the liver 48 hours after PHx. (G) Statistical analysis of the number of Ki-67-expressing cells after PHx (n=3). The number of Ki-67-positive cells was normalized to the number of DAPI-stained dots. Scale bar = 100 µm. Student *t*-test; \**p* < 0.05, \*\**p* < 0.01, \*\*\**p* < 0.001. Data are expressed as the mean ± SEM.

To investigate hepatocyte proliferation between *Prom1^f/f^* and *Prom1^LKO^* mice during liver regeneration, we confirmed cell cycle-related genes (Cyclin A, B, D, E, and PCNA) in PHx livers by qRT-PCR and immunoblotting. The levels of cyclin mRNAs were reduced in *Prom1^LKO^* livers more than in *Prom1^f/f^* livers (Fig. 2B). Consistently, the expression of cell cycle-related proteins in *Prom1^LKO^* mice decreased compared to that in *Prom1^f/f^* mice after PHx (Fig. 2C, D). We also analyzed hepatocyte proliferation by H&E staining and double immunofluorescence along with Ki-67 (as a cell proliferation marker) and HNF4α (Fig. 2E-G). As shown in Fig. 2F, Ki-67 expression in *Prom1^f/f^* livers increased more than that in *Prom1^LKO^* livers after PHx. Indeed, PROM1 deficiency reduced the number of Ki-67-positive cells by ∼50% (Fig. 2G). These results suggested that the liver-specific deletion of PROM1 decreased hepatocyte proliferation and impaired liver regeneration after PHx.

### Liver-specific PROM1 deficiency reduces liver regeneration in mice injected with CCl_4_

To further investigate the effects of PROM1 deficiency on the proliferation of hepatocytes in the regenerating liver, we analyzed the liver after injecting CCl_4_ into mice. As with liver regeneration by PHx, PROM1 expression also increased after CCl_4_ injection. PROM1 mRNA increased over ∼10-fold in the liver by CCl_4_ (Fig. 3A). PROM1 double immunofluorescence with HNF4α or CK19 showed that major cells expressing PROM1 were hepatocytes after CCl_4_ injection (Fig. 3B, C).

**Figure 3.**
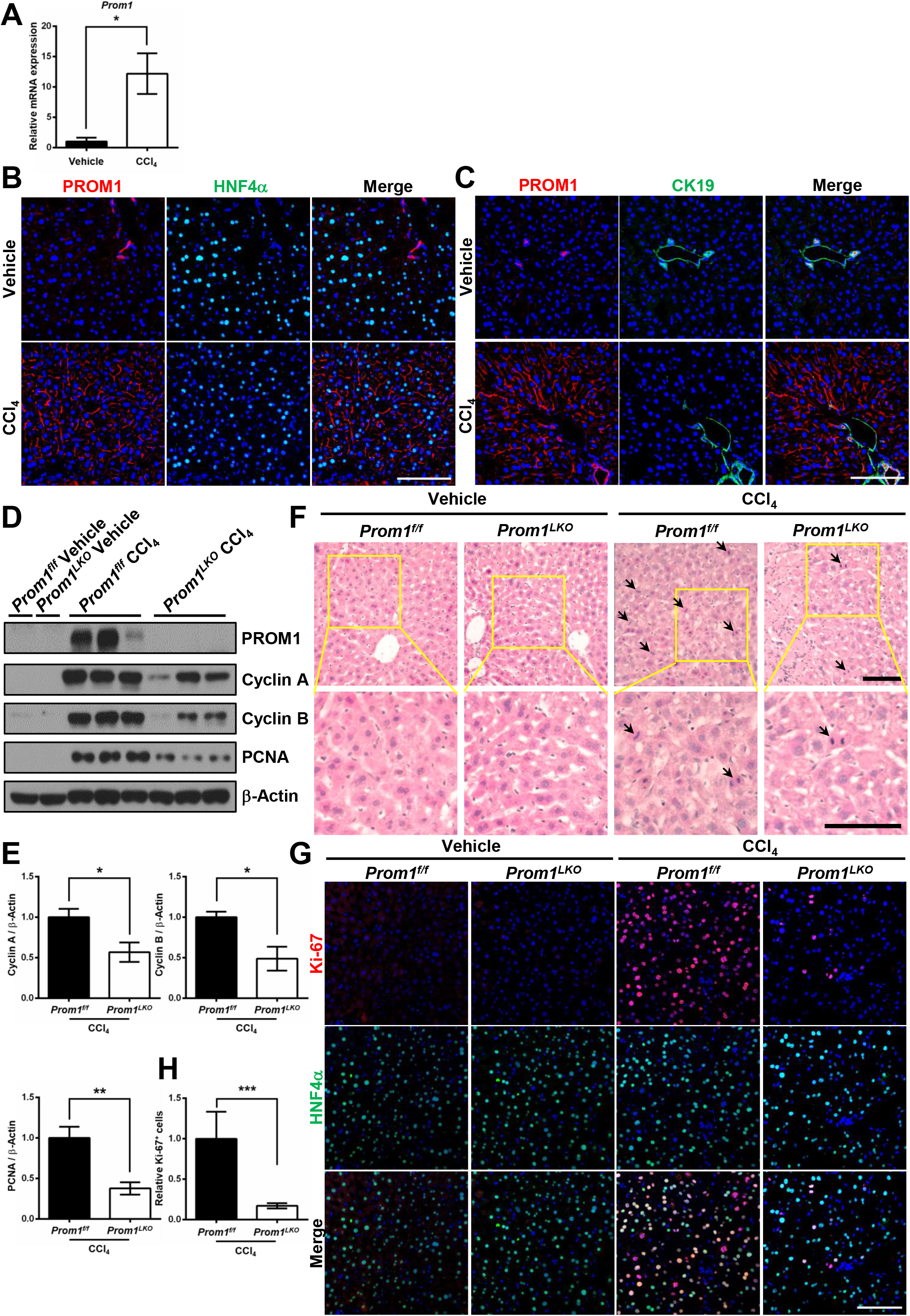
Liver-specific deletion of *Prom1* in mice impairs liver regeneration after CCl_4_ injection. Eight-week-old male *Prom1^f/f^* and *Prom1^LKO^* mice were intraperitoneally injected with vehicle (n=3) or CCl_4_ (n=5) for 48 hours. The liver was analyzed by qRT-PCR, immunoblotting and immunofluorescence. (A) The relative mRNA level for PROM1. The mRNA level of PROM1 was normalized by 18S rRNA. (B, C) Double immunofluorescence for PROM1 and ΗNF4α (B) or CK19 (C). (D) Immunoblotting for PROM1, Cyclin A and B, and PCNA. (E) Statistical analysis of the band intensity in D. The band intensity of each protein was normalized to that of β-actin. (F) Representative H&E staining in the liver. Mitotic cells are indicated by arrows. (G) Double immunofluorescence for Ki-67 and HNF4α. (H) Statistical analysis of the number of Ki-67- expressing cells (n=3). The number of Ki-67-positive cells was normalized to the number of DAPI- stained dots. Scale bar = 100 µm. Student *t*-test; \**p* < 0.05, \*\**p* < 0.01, \*\*\**p* < 0.001. Data are expressed as the mean ± SEM.

Next, we compared the expression of cell cycle-related proteins in the livers of *Prom1^f/f^* and *Prom1^LKO^* mice after CCl_4_ injection. PROM1 deficiency significantly decreased the expression of Cyclin A, Cyclin B, and PCNA, as determined by immunoblotting (Fig. 3D, E). Hepatocyte proliferation was confirmed by H&E staining in the livers of *Prom1^f/f^* and *Prom1^LKO^* mice after CCl_4_ injection (Fig. 3F). PROM1 deficiency decreased the number of Ki-67-expressing cells after CCl_4_ injection by ∼80%, as determined by immunofluorescence (Fig. 3G, H). Taken together, these data suggested that PROM1 deficiency attenuates hepatocyte proliferation during liver regeneration in the CCl_4_ model.

### PROM1 increases IL-6 signaling during liver regeneration

Hepatocyte proliferation in the early stage of liver regeneration requires the JAK-STAT, PI3K, MAPK, and β-catenin signaling pathways initiated by different mitogens, such as IL-6, EGF, HGF and Wnt [2, 17, 18]. To examine the signaling pathways affected by PROM1, we observed the expression and activation of these mitogenic signaling molecules after PHx by immunoblotting. PROM1 deficiency significantly decreased the phosphorylation status of STAT3 and ERK but not the phosphorylation status of AKT or GSK3β (Fig. 4 A, B). In the CCl_4_ model, PROM1 deficiency also decreased the phosphorylation of STAT3 (Fig. 4 C, D). Since IL-6 signals are known to activate both STAT3 and ERK, these results led us to investigate the IL-6 signaling pathway in more detail. IL-6 ELISA showed that PROM1 deficiency did not change the serum level of IL-6 after PHx (Fig. 4E), thus allowing us to rule out the effect of PROM1 on IL-6 production and secretion during liver regeneration. Next, we confirmed that PROM1 overexpression statistically increased IL-6-induced phosphorylation of STAT3 by ∼2-fold in HEK 293 cells and by ∼6-fold in primary hepatocytes obtained from *Prom1^LKO^* mice (Fig. 4F, G).

**Figure 4.**
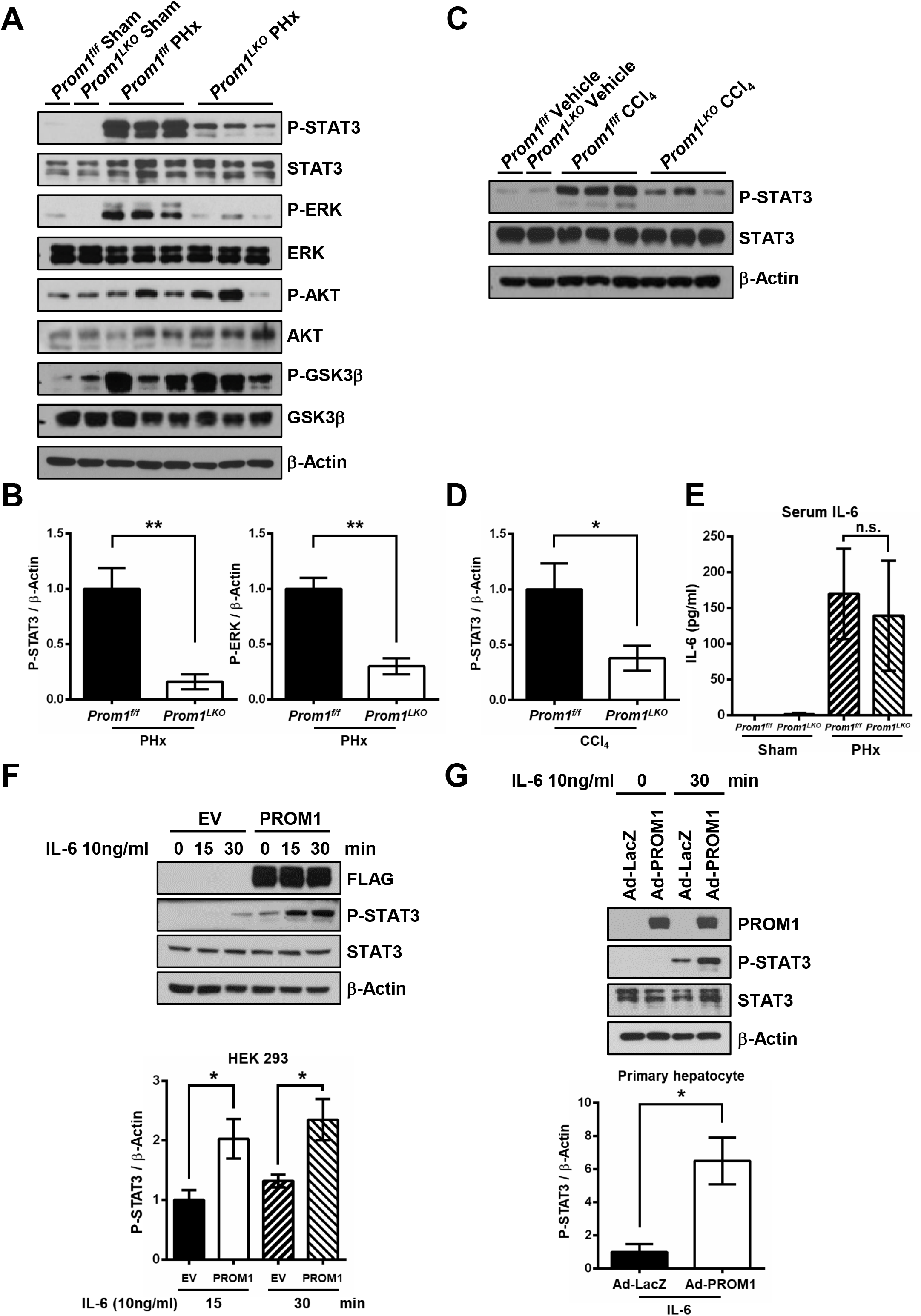
PROM1 facilitates IL-6-STAT3 signaling pathway. (A, B) A 2/3 partial hepatectomy was performed in 8-week-old male *Prom1^f/f^* and *Prom1^LKO^* mice. The liver was analyzed by immunoblotting 24 hours after PHx (n=5-6). Immunoblotting for STAT3, P-STAT3, ERK, P-ERK, AKT, P-AKT, GSK3β, and P-GSK3β (A). Statistical analysis of the band intensity of P-STAT3 and P-ERK. The band intensity of each protein was normalized to that of β-actin (B). (C, D) Eight-week-old male *Prom1^f/f^* and *Prom1^LKO^* mice were intraperitoneally injected with vehicle (n=3) or CCl_4_ (n=5) for 48 hours. Immunoblotting for STAT3, and P-STAT3 (C). Statistical analysis of the band intensity of P-STAT3. The band intensity of each protein was normalized to that of β-actin (D). (E) Quantification of serum IL-6 in *Prom1^f/f^* and *Prom1^LKO^* mice (n=4-6) 24 hours after PHx. (F, G) Empty vector (EV) or FLAG-tagged PROM1 was transfected into HEK293 cells for 48 hours. After serum starvation for 16 hours, cells were treated with 10 ng/ml human recombinant IL-6 for 0, 15, and 30 minutes (F). Primary hepatocytes were isolated from 8-week-old male *Prom1^LKO^* mice. The cells were infected with adeno-LacZ or PROM1 for 16 hours, followed by serum starvation for 16 hours and then harvested after treatment with 10 ng/ml IL-6 for 30 minutes (G). Each experiment was independently repeated three times. Immunoblotting for FLAG or PROM1, STAT3 and P-STAT3. Statistical analysis of the band intensity of P-STAT3. The intensity of P-STAT3 was normalized to that of β-actin. Student *t*-test; \**p* < 0.05, \*\**p* < 0.01, n.s., nonsignificant. Data are expressed as the mean ± SEM.

To further confirm the association between PROM1 and the IL-6 signaling pathway, we observed whether liver regeneration impaired by PROM1 deficiency was rescued through adenoviral overexpression of constitutively activated STAT3 (Stat3c) in *Prom1^LKO^* mice. As determined by qRT-PCR and immunoblotting (Fig. S1A-C), cyclins A, B, and E and PCNA were significantly increased by Stat3c overexpression. Consistent with these data, Ki-67 and HNF4α immunofluorescence showed that hepatocyte proliferation was increased by Stat3c in *Prom1^LKO^* mice because the number of Ki-67-expressing cells increased by ∼4-fold (Fig. S1D, E). These results suggested that PROM1 regulates hepatocyte proliferation through the IL-6 signaling pathway during liver regeneration.

### PROM1 regulates IL-6 signaling by interacting with GP130 in lipid rafts

GP130, a common receptor of the IL-6 receptor family and known as the IL-6 receptor beta-subunit signal transducer, associates with downstream molecules in lipid rafts for efficient signaling [19, 20]. Since both PROM1 and IL-6 signaling complexes were in lipid rafts, we hypothesized that PROM1 would bind to GP130. To prove this hypothesis, we investigated whether raft localization of GP130 is dependent on PROM1 in PHx livers. We confirmed that PROM1 deficiency reduced the expression of GP130 in liver lipid rafts (Fig. 5A). In addition, PROM1 overexpression in *Prom1^LKO^* sham livers increased the expression of GP130 in liver lipid rafts (Fig. 5B).

**Figure 5.**
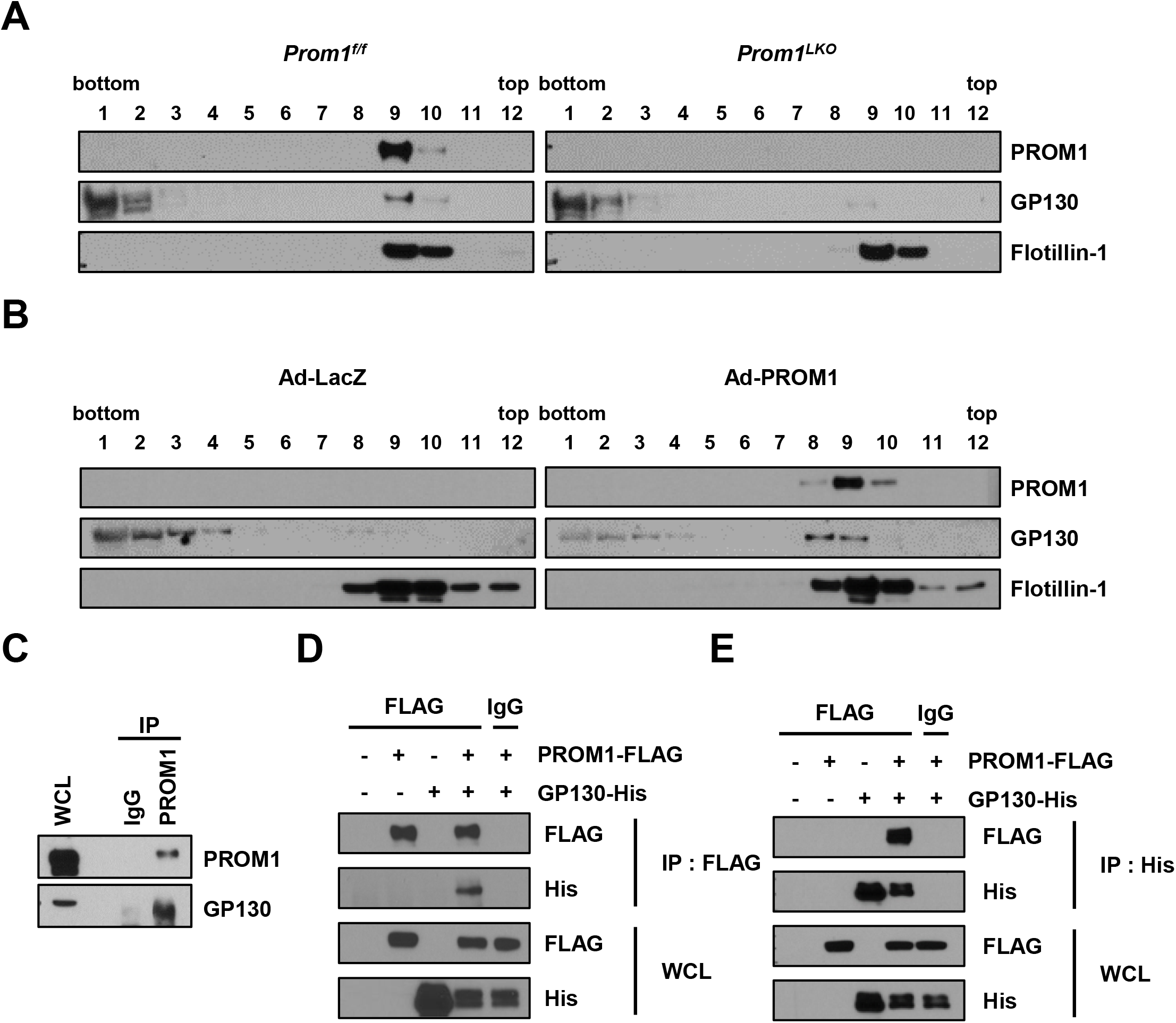
PROM1 interacts with GP130 in lipid rafts. (A) Detergent-resistant lipid rafts were isolated from *Prom1^f/f^* and *Prom1^LKO^* mouse livers 48 hours after PHx. Protein expression levels of PROM1, GP130, and Flotillin-1 were determined by immunoblotting in each fraction after sucrose gradient ultracentrifugation. (B) Detergent-resistant lipid rafts were isolated from 8-week-old male *Prom1^LKO^* mice 3 days after infection with adeno-LacZ or adeno-PROM1. Immunoblotting for PROM1, GP130, and Flotillin-1 in each fraction. (C) Coimmunoprecipitation was performed with normal IgG or anti-PROM1 in wild-type livers 48 h after PHx. Immunoblotting for endogenous PROM1 and GP130. (D, E) The molecular interaction between PROM1 and GP130 was determined by reciprocal immunoprecipitation after PROM1-FLAG and GP130-His were transfected into HEK 293 cells for 48 hours. WCL, whole cell lysates; IP, immunoprecipitation; IgG, normal IgG.

Next, we demonstrated the molecular interaction between PROM1 and GP130 by immunoprecipitation. As shown in Fig. 5C-E, endogenous immunoprecipitation in PHx wild-type liver and reciprocal exogenous immunoprecipitation in HEK 293 cells showed a molecular interaction between PROM1 and GP130. All these data indicate that PROM1 binds to GP130, which is important for the raft localization of GP130.

### The first extracellular domain of PROM1 is required for the interaction with GP130 and the regulation of the STAT3 signaling pathway

To determine the domain required for the interaction between PROM1 and GP130, we generated various deletion mutants of PROM1 (Fig. 6A). A coimmunoprecipitation assay using these mutants showed that all deletion mutants of PROM1 still interacted with GP130 (Fig. 6B). Based on this result, we hypothesized that the first extracellular domain of PROM1 (PROM1-EX1) would be an important region for the interaction between the two proteins. To prove this hypothesis, we generated GPI-anchored PROM1-EX1 (PROM1^GPI-EX1^) in which the first transmembrane domain was substituted with a GPI anchor and observed the interaction between PROM1^GPI-EX1^ and GP130. As shown Fig. 6C, EX1 itself interacted with GP130.

**Figure 6.**
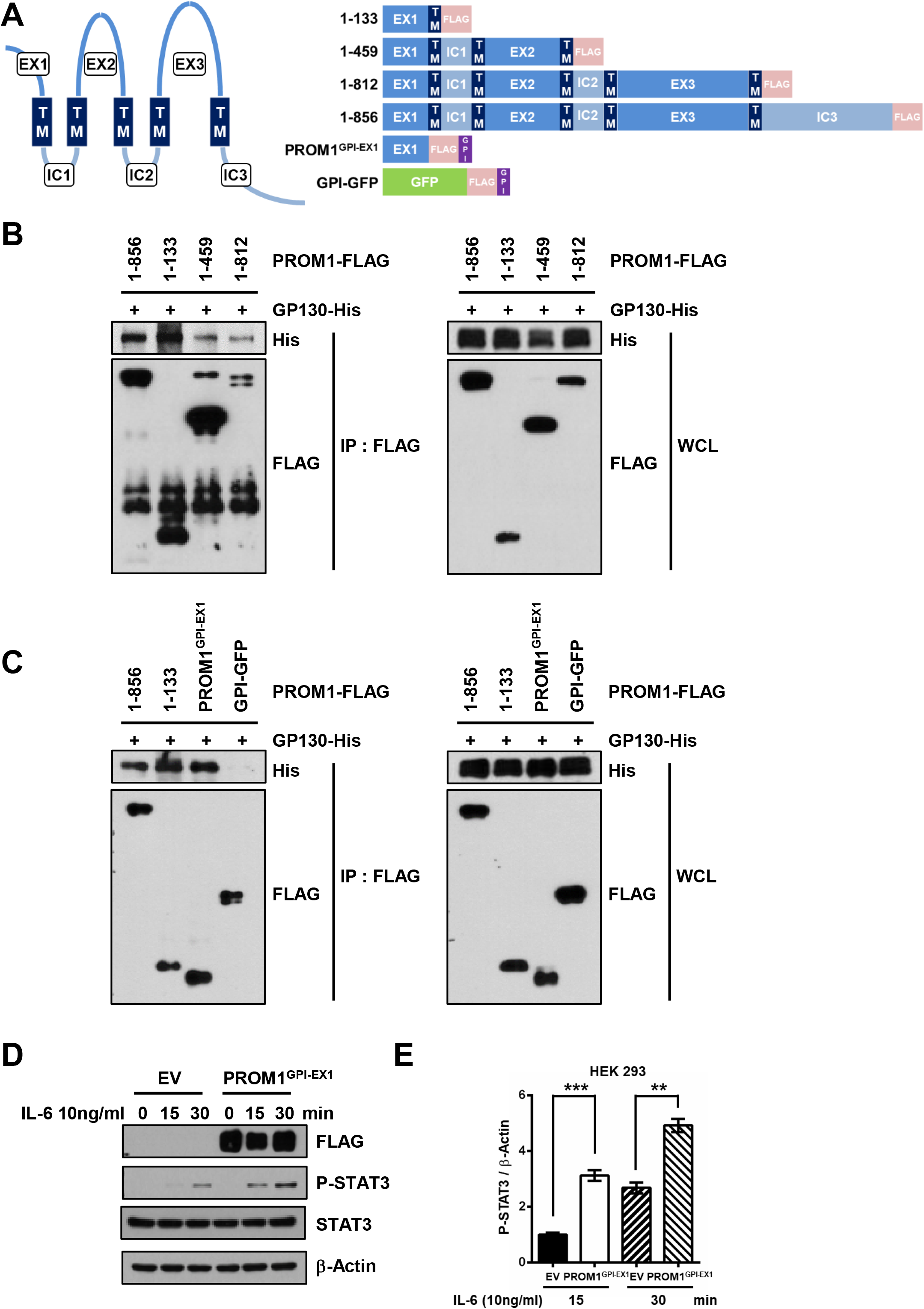
The first extracellular domain of PROM1 interacts with GP130 and regulates the STAT3 signaling pathway. (A) Structures of PROM1 deletion mutants. EX, extracellular domain; TM, transmembrane domain; GPI, glycosylphosphatidylinositol; GFP, green fluorescence protein. (B, C) Coimmunoprecipitation between each PROM1 mutant and GP130. HEK 293 cells were transfected with various FLAG-tagged PROM1 mutants (1-133, 1-459, 1-812, and PROM1^GPI-EX1^) or full-length PROM1 (1-856) and His-tagged GP130 for 48 hours. (D, E) HEK 293 cells were transfected with empty vector (EV) or FLAG-tagged PROM1^GPI-EX1^ for 48 hours. After serum starvation for 16 hours, HEK 293 cells were treated with 10 ng/ml IL-6 for 0, 15, or 30 minutes. The experiments were independently repeated three times. Immunoblotting for STAT3, P-STAT3 and FLAG (D). Statistical analysis of the band intensity of P-STAT3. The band intensity of P- STAT3 was normalized to that of β-actin (E). WCL, whole cell lysates; IP, immunoprecipitation; IgG, normal IgG. Student *t*-test; \*\**p* < 0.01, \*\*\**p* < 0.001. Data are expressed as the mean ± SEM.

Since GPI-anchored proteins are expressed in lipid rafts, we examined whether PROM1^GPI-EX1^ enhances the STAT3 signaling pathway. Exogenous PROM1^GPI-EX1^ itself increased the activity of STAT3 in HEK 293 cells (Fig. 6D, E). Taken together, the first extracellular domain of PROM1 is required for binding to GP130 and regulating the GP130-STAT3 signaling pathway.

### The expression of GPI-anchored PROM1-EX1 rescues liver regeneration in PROM1-deficient mice after partial hepatectomy

To evaluate whether PROM1^GPI-EX1^ has an in vivo function in liver regeneration after PHx, we observed recovery of liver mass and hepatocyte proliferation after adenoviral overexpression of PROM1^GPI-EX1^ in *Prom1^LKO^* mice. PROM1^GPI-EX1^ was overexpressed in the liver, as determined by qRT-PCR and immunoblotting (Fig. 7B, C). The overexpression of PROM1^GPI-EX1^ alone was sufficient to increase the liver-to-body weight ratio at 24 and 48 hours after PHx in *Prom1^LKO^* mice (Fig. 7A). As determined by qRT-PCR and/or immunoblotting for Cyclin A and B, PCNA and phospho-STAT3 and immunofluorescence for Ki-67, the overexpression of PROM1^GPI-EX1^ statistically increased hepatocyte proliferation via STAT3 phosphorylation compared to the overexpression of LacZ (Fig. 7B-H). In addition, GP130 was relocalized into lipid rafts after the overexpression of PROM1^GPI-EX1^ (Fig. 7I). These data suggested that PROM1^GPI-EX1^ has a crucial role in refining GP130 into lipid rafts and mediating an IL-6-GP130 axis, thereby promoting liver regeneration.

**Figure 7.**
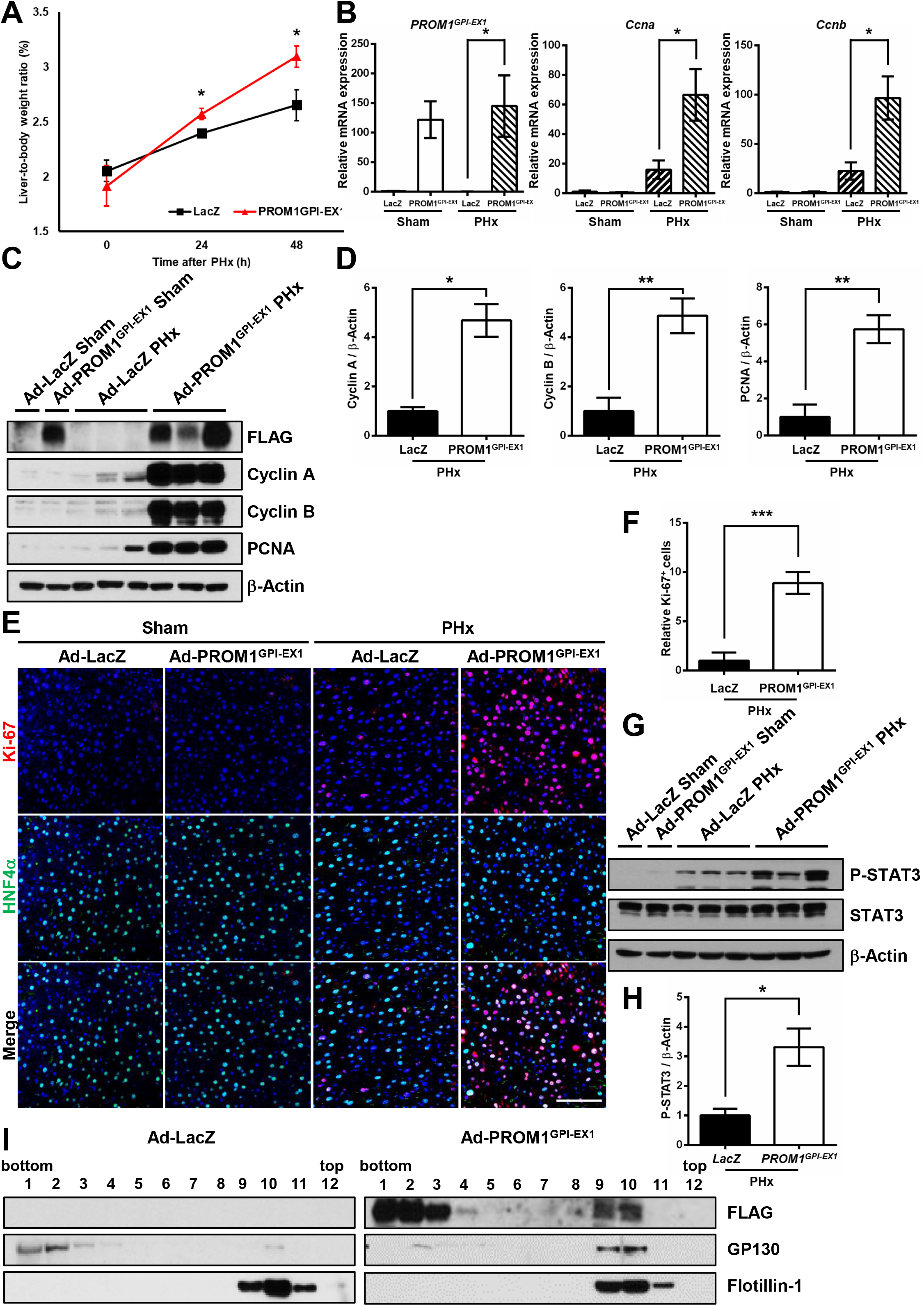
The expression of GPI-anchored PROM1-EX1 rescues liver regeneration in liver-specific *Prom1*-deficient mice. A 2/3 partial hepatectomy was performed in 8-week-old male *Prom1^LKO^* mice after infection with adeno-LacZ (n=3-6) or adeno-PROM1^GPI-EX1^-FLAG (n=4-9). (A) Ratio of liver-to-body weight on the indicated days after PHx. (B) The relative mRNA levels of *PROM1^GPI-EX1^*, *Ccna*, and *Ccnb* in the liver 24 hours after PHx. Each mRNA level was normalized by 18S rRNA. (C, D) Immunoblotting for Cyclin A and B, PCNA, and FLAG in the liver 48 hours after PHx (C). Statistical analysis of the band intensities of Cyclin A and B and PCNA in C. The band intensity of each protein was normalized to that of β-actin (D). (E, F) Double immunofluorescence for Ki-67 and HNF4α in the liver 48 hours after PHx (E). Statistical analysis of the number of Ki-67-expressing cells (n=3). The number of Ki-67-positive cells was normalized to the number of DAPI-stained dots (F). (G, H) Immunoblotting for STAT3, and P-STAT3 in the liver 24 hours after PHx (G). Statistical analysis of the band intensity of P-STAT3 in G. The band intensity of P-STAT3 was normalized to that of β-actin (H). (I) Detergent-resistant lipid rafts were isolated from adeno-LacZ or adeno-PROM1^GPI-EX1^-FLAG mouse livers 48 hours after PHx. Protein expression levels of FLAG, GP130, and Flotillin-1 were determined by immunoblotting in each fraction after sucrose gradient ultracentrifugation. Scale bar = 100 µm. Student *t*-test; \**p* < 0.05, *\*\*p* < 0.01, \*\*\**p* < 0.001. Data are expressed as the mean ± SEM.

## Discussion

PROM1 is well known as a marker for cancer stem cells and normal stem cells. Recent studies have revealed its ability to regulate various cellular signal transduction pathways by interacting with PI3K, HDAC6, radixin, and SMAD7 [15, 16, 21, 22]. Here, we demonstrated that PROM1 is also necessary for regulating IL-6 signaling during liver regeneration. We found that the expression of PROM1 dramatically increased in hepatocytes during liver regeneration after PHx or CCl_4_ injection. Hepatocellular PROM1 facilitated the IL-6 signaling pathway by interacting with GP130 in lipid rafts. As a result, we demonstrated that PROM1 promoted the proliferation of hepatocytes during liver regeneration (Fig. 8).

**Figure 8.**
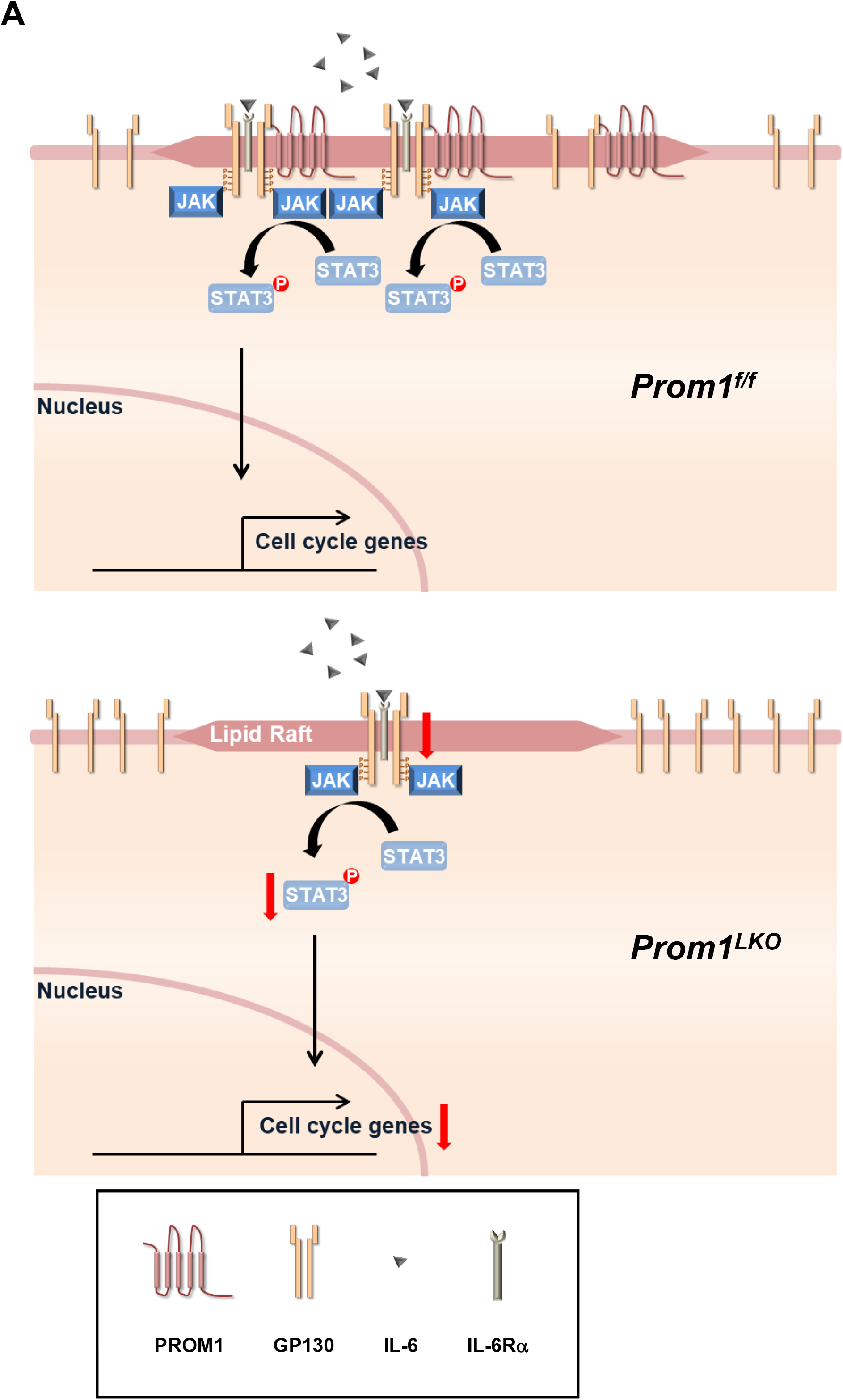
PROM1 promotes hepatocyte proliferation through facilitating IL-6-GP130 signaling pathway in lipid rafts during liver regeneration. (A) After 2/3 partial hepatectomy or CCl_4_ injection, PROM1 expression increases in hepatocytes. Upregulated PROM1 recruits GP130 into lipid rafts by interacting with GP130. PROM1/GP130 complex in lipid rafts facilitates the IL- 6-GP130-STAT3 signaling pathway. Therefore, PROM1 promotes hepatocyte proliferation during liver regeneration.

During liver regeneration after PHx and CCl_4_ injection, PROM1 was highly upregulated, as determined by qRT-PCR, immunofluorescence, and immunoblotting. In addition, PROM1 upregulation was dramatically demonstrated in PROM1 lineage tracing mice (*Prom1^Cre/ERT2^; Rosa26^tdTomato^* mice), in which cells express tdTom under the control of the PROM1 promoter. Because hepatocellular PROM1 upregulation is also observed after bile duct ligation (BDL) [16] and a lithogenic diet (data not shown), various liver damages might lead to hepatocellular PROM1 upregulation. Many extracellular and intracellular factors, such as HIF-1α, TGFβ1, p53 and mTOR, regulate the expression of PROM1 [12, 23]. A previous study reported that STAT3 promotes the transcription of PROM1 in hepatocellular carcinoma [24]. Therefore, we hypothesized that the IL-6-STAT3 signaling pathway might be necessary for upregulating PROM1 in hepatocytes, and then, the upregulated PROM1 would form a ‘positive loop’ because PROM1 promotes the IL-6-STAT3 signaling pathway.

PROM1 interacts with various signaling molecules through its different domains. The cytoplasmic C-terminal domain of PROM1 binds to PI3K and radixin, maintaining cancer stem cell properties and regulating glucagon-induced PKA activity, respectively [15, 21]. The first intracellular loop of PROM1 binds to HDAC6 and SMAD7, regulating β-Catenin signaling and TGFβ signaling, respectively [16, 22]. Here, we demonstrated that the first extracellular domain of PROM1 binds to GP130. Furthermore, lipid raft-targeted PROM1^GPI-EX1^ alone is sufficient to replace the function of full-length PROM1, which recruits GP130 into lipid rafts and then facilitates IL-6-induced STAT3 phosphorylation, leading to hepatocyte proliferation and liver regeneration.

The PROM1-positive population in various tumors has self-renewal and differentiation potential and chemotherapy or radiotherapy resistance [11]. Although most cancers are removed through cancer therapy, only a small number of surviving PROM1-positive cells can proliferate and cause cancer to recur. Thus, PROM1 has been considered a very important target protein for cancer therapy. Because PROM1 expression was upregulated at the early stage of liver regeneration (within 48 hours after PHx) and then returned to sham liver levels at the termination stage of regeneration (7 days after PHx, data not shown), hyperplasia or tumorigenesis might not occur during liver regeneration.

In addition to IL-6, GP130 is involved in various signaling pathways of IL-6 family cytokines, such as IL-11, leukemia inhibitory factor (LIF), oncostatin M, and ciliary neurotrophic factor [25]. Therefore, the PROM1-GP130 axis could be a potential therapeutic target for human diseases induced by these cytokines. For example, a PROM1-neutralizing antibody targeting PROM1-EX1 is a good candidate for alleviating inflammatory diseases caused by these cytokines.

## Materials and methods

### Animal studies

Whole-body *Prom1* knockout mice (*Prom1^Cre/ERT2-nlacZ^*) were purchased from The Jackson Laboratory (Stock No: 017743, Bar Harbor, ME, USA) and backcrossed with C57BL/6N mice for five generations. Liver-specific *Prom1* knockout mice were generated by crossing *Prom1 ^flox/flox^* C57BL/6 mice (ToolGen, Seoul, Korea) with *Alb-Cre* C57BL/6 mice containing the Cre recombinase sequence driven by the albumin promoter (The Jackson Laboratory, Bar Harbor, ME, USA). *Prom1* lineage tracing mice were generated by crossing *Prom1^Cre/ERT2-nlacZ^* C57BL/6 mice with *Rosa26^tdTomato^* C57BL/6 mice containing the tdTomato sequence prevented by the loxP-flanked STOP cassette (Stock No: 007914, The Jackson Laboratory, Bar Harbor, ME, USA). For Cre-loxP recombination, tamoxifen (T5648; Sigma, 20 mg/ml in corn oil) was intraperitoneally injected at 150 mg/kg 1 day before 2/3 partial hepatectomy in 8-week-old mice.

All mice were housed in plastic cages under a 12:12-hour light/dark photoperiod at controlled temperature with free access to water and food. All mice were bred, maintained, and cared for in a manner consistent with criteria outlined in the Principles of Laboratory Animal Care (NIH publication no. 85-23, revised 1985). Protocols for animal studies were approved by the Institutional Animal Care and Use Committee of Korea University and the Korean Animal Protection Law (KUIACUC-2019-0111).

To investigate liver regeneration in mice, a 2/3 partial hepatectomy and CCl_4_ injection were performed. For the 2/3 partial hepatectomy model, two-thirds of the mouse liver was surgically removed as previously described [26, 27]. Briefly, 8-week-old mice were anesthetized using isoflurane. An abdominal midline incision was made to open the abdominal cavity to expose the liver. The hepatic left lateral and median lobes were isolated and ligated. After ligation, each lobe was removed with surgical scissors. Then, the abdominal skin was sutured and sterilized. After surgery, the mice were kept warm for recovery. For the CCl_4_ model, 8-week-old mice were intraperitoneally injected with 25% CCl_4_ in corn oil (Sigma) at a dose of 2.4 µl/g body weight.

### Mouse primary hepatocytes isolation

Primary hepatocyte isolation was performed based on two-step collagenase perfusion as previously described [15]. Briefly, 8-week-old mice were anesthetized with avertin (intraperitoneal injection of 250 mg/kg body weight). After an abdominal midline incision, the livers were perfused with EGTA-containing perfusion buffer (140 mM NaCl, 6 mM KCl, 10 mM HEPES, and 0.08 mg/mL EGTA, pH 7.4) at a rate of 7 ml/min for 5 min, followed by continuous perfusion with collagenase-containing buffer (66.7 mM NaCl, 6.7 mM KCl, 5 mM HEPES, 0.48 mM CaCl_2_, and 3 g/mL type IV collagenase, pH 7.4) for 8 min. After collecting parenchymal cells by low-speed (50 × g, 4 min) centrifugation, viable hepatocytes were purified by Percoll gradient centrifugation. Then, hepatocytes were resuspended in complete growth medium (M199 media containing 10% fetal bovine serum, 23 mM HEPES, and 10 nM dexamethasone) and seeded on collagen-coated plates at a density of 3.3 × 10^5^ cells/ml. After 4 hours of cell attachment, the medium was replaced with complete growth medium and replaced daily before use in all experiments. For in vitro analysis of IL-6-induced STAT3 phosphorylation, cells were treated with human recombinant interleukin-6 (Peprotech).

### Adenovirus preparation and infection

Adenoviruses harboring LacZ, PROM1, STAT3C (#99264, Addgene), and PROM1^GPI-EX1^ were prepared as previously described [28]. AD293 cells were infected with each viral stock to amplify the viruses. Virus purification was performed by double cesium chloride-gradient ultracentrifugation. Viral particles in cesium chloride (density≒1.345) were collected and washed with washing buffer (10 mM Tris pH 8.0, 2 mM MgCl_2_, and 5% sucrose). Purified adenoviruses (0.5 × 10^9^ pfu) were intravenously injected into the tails of mice.

### RNA isolation and quantitative RT–PCR

Total RNA was extracted from liver tissues using an easy-spin^TM^ total RNA extraction kit (Intron Biotechnology, Korea) according to the manufacturer’s protocol. Total RNA (4 µg) was used for cDNA synthesis using random hexamers, oligo dT primers, and reverse transcription master mix (ELPIS Biotech, Korea). Quantitative real-time PCR was performed using the cDNAs and each gene-specific oligonucleotide primer in the presence of TOPreal qPCR premix (Enzynomics, Korea). The following real-time PCR conditions were used: an initial denaturation step at 95 °C for 15 min, followed by 50 cycles of denaturation at 95 °C for 10 sec, annealing at 58 °C for 15 sec, and elongation at 72 °C for 20 sec. Each PCR product was evaluated by melting curve analysis for quality control. Supplementary Table 1 shows the sequences of the gene-specific primers used for qRT-PCR.

### Immunoblotting and Immunoprecipitation

To extract whole cell lysates, the livers were homogenized with a tissue homogenizer and harvested. The homogenized tissues were lysed with radioimmunoprecipitation assay buffer (50 mM Tris-Cl pH 8.0, 150 mM NaCl, 1% NP-40, 0.5% sodium deoxycholate, 0.1% sodium dodecyl sulfate, and protease- and phosphatase-inhibitor cocktail (Gendepot, USA)) on ice for 30 min. Whole cell lysates were extracted from supernatant by microcentrifugation at 14,000 rpm for 10 min at 4 °C. The whole cell lysates were quantified by BCA assay. The normalized protein samples were separated by sodium dodecyl sulfate–polyacrylamide gel electrophoresis. The separated proteins were transferred to a nitrocellulose membrane and incubated with the primary antibodies of interest (Supplementary Table 2) followed by incubation with horseradish peroxidase (HRP)- conjugated secondary antibodies. The protein band signals were visualized by chemiluminescence detection using an EZ-Western kit (Dogenbio, Korea).

For immunoprecipitation, homogenized tissues or cells were lysed with buffer containing 25 mM HEPES, 150 mM NaCl, 1% NP-40, 10 mM MgCl_2_, 1 mM EDTA, 2% glycerol, and protease inhibitor cocktail (Gendepot, USA) on ice for 30 min. Whole cell lysates were extracted from supernatant by microcentrifugation at 14,000 rpm for 10 min at 4 °C. The whole cell lysates were quantified by BCA assay. One milligram of protein in whole cell lysates was incubated with specific primary antibody overnight at 4 °C, followed by incubation with 60 µg of Protein A- or G-agarose bead slurry (Roche, Germany) for 4 hours at 4 °C. The bead precipitates were washed with buffer containing 25 mM HEPES, 150 mM NaCl, 1% NP-40, 10 mM MgCl_2_, 1 mM EDTA, 2% glycerol, and protease inhibitor cocktail (Gendepot, USA) 4 times. Protein samples were obtained from the precipitates and analyzed by immunoblotting as described above.

### Immunofluorescence staining

For immunofluorescence staining of liver tissues, frozen tissues were cut to a thickness of 5 µm using a cryocut microtome (Leica).

For PROM1 double immunofluorescence with HNF4α or CK19, the sections were incubated with proteinase K (0.06 U/mg) for 5 min, followed by blocking with 2.5% normal horse serum for 30 min at room temperature. Then, the sections were incubated with mouse anti-HNF4α (Abcam) or rabbit anti-CK19 (Abcam) overnight at 4 °C. Next, for double immunofluorescence with PROM1, the sections were incubated with 4% paraformaldehyde in 0.1 M phosphate buffer for 30 min at 37 °C and then incubated with rat anti-PROM1 (Thermo Fisher Scientific) overnight at 4 °C. Then, the sections were incubated with fluorescence-conjugated secondary antibody (Thermo Fisher Scientific) for 1 hour at room temperature.

For tdTom or Ki-67 double immunofluorescence with HNF4α or CK19, heat-mediated antigen retrieval using a pressure cooker in citrate buffer (pH 6.0) was performed on frozen sections. After antigen retrieval, the sections were blocked with 2.5% normal horse serum for 30 min at room temperature. Then, the sections were incubated with rabbit (Rockland) or rat (Thermo Fisher Scientific) anti-tdTom, rabbit anti-Ki-67 (Cell Signaling Technology), mouse anti-HNF4α (Abcam), and rabbit anti-CK19 (Abcam) overnight at 4 °C and then incubated with fluorescence-conjugated secondary antibody (Thermo Fisher Scientific) for 1 hour at room temperature. After mounting with Fluoroshield^TM^ with DAPI (Sigma), the images were captured using an LSM800 confocal microscope (Zeiss).

### Immunohistochemistry

For hematoxylin-eosin staining of liver tissues, paraffin-embedded tissues were cut to a thickness of 5 µm using a multirotary microtome (Leica). The sections were stained with hematoxylin-eosin according to a standard protocol. After mounting with synthetic mountant (Thermo Fisher Scientific), the images were captured using a light microscope (Leica).

### Serum IL-6 ELISA

Serum IL-6 levels were quantified using a commercial mouse IL-6 ELISA kit (RAB0308, Sigma) according to the manufacturer’s instructions and analyzed using spectra-iMAX (Molecular Devices).

### Detergent-resistant lipid rafts isolation

To obtain detergent-resistant lipid rafts, homogenized liver tissues were lysed with buffer containing 1% Brij-35, 25 mM HEPES pH 6.5, 150 mM NaCl, 1 mM EDTA, and protease inhibitor cocktail (Gendepot, USA) on ice for 30 min. Then, the lysates were subjected to discontinuous sucrose gradient (40, 35, and 5%) ultracentrifugation using a SW41Ti rotor (28,7000 × g) for 18 hours at 4 °C. After ultracentrifugation, the sucrose solutions were fractionated into 12 fractions. A cloudy band corresponding to the lipid rafts was collected at the interface between the 35 and 5% sucrose solutions and confirmed by immunoblotting for Flotillin-1 as a lipid raft marker.

### Plasmid construction and transient transfection

Deletion mutants of FLAG-tagged human PROM1 transcript variant 2 (PROM1-FLAG) were generated by reverse PCR as previously described [15]. FLAG-tagged GPI-anchored PROM1- EX1 was generated by the DNA assembly method (#E2621, NEB, Ipswich, MA, USA). The GPI-anchor signal sequence from pCAG:GPI-GFP (#32601, Addgene) was added at the C-terminus of PROM1-EX1 (1-99)-FLAG. His-tagged GP130 was generated by adding a 6×His tag sequence at the C-terminus of the GP130 CDS obtained from the cDNA library of HEK293 cells.

DNA transfection was performed using Lipofectamine 3000 reagent (Invitrogen, USA) according to the manufacturer’s instructions.

### Statistical analysis

The number of mice used in each experiment was determined based on preliminary experiments in the same model. Immunofluorescence images and immunoblotting band intensities were quantified using ImageJ (NIH) or Photoshop (Adobe) software. The images used for statistics contained more than ∼250 cells per field and were taken from a minimum of 3∼5 fields per sample. Data are expressed as the mean ± SEM. Sample numbers are indicated in the figure legends. A two-tailed Student’s *t*-test was used to calculate the *p value*s. Significance levels were \**p* < 0.05; \*\**p* < 0.01; \*\*\**p* < 0.001; and n.s., nonsignificant. A *p value* < 0.05 was considered statistically significant.

## Author contributions

M.-S.B., D.-M.Y., M.L., S.-J.J., J.-W.L., H.L., A.K., and J.S.K. performed the experiments; J.- H.H.,S.-H.K., J.-S.L. and Y.-G.K. designed the experiments and analyzed the data; and M.-S.B., D.-M.Y. and Y.-G.K. wrote the manuscript.

## Acknowledgement

We thank all members of our laboratory for their supports and intellectual inputs during the preparation of this manuscript. Funding: This work was supported by grants from the National Research Foundation of Korea awarded to; Y.-G. Ko (R1509597 and R20000552)

## Conflicts of interest

The authors disclose no conflicts

---

**Supplementary Figure 1.**
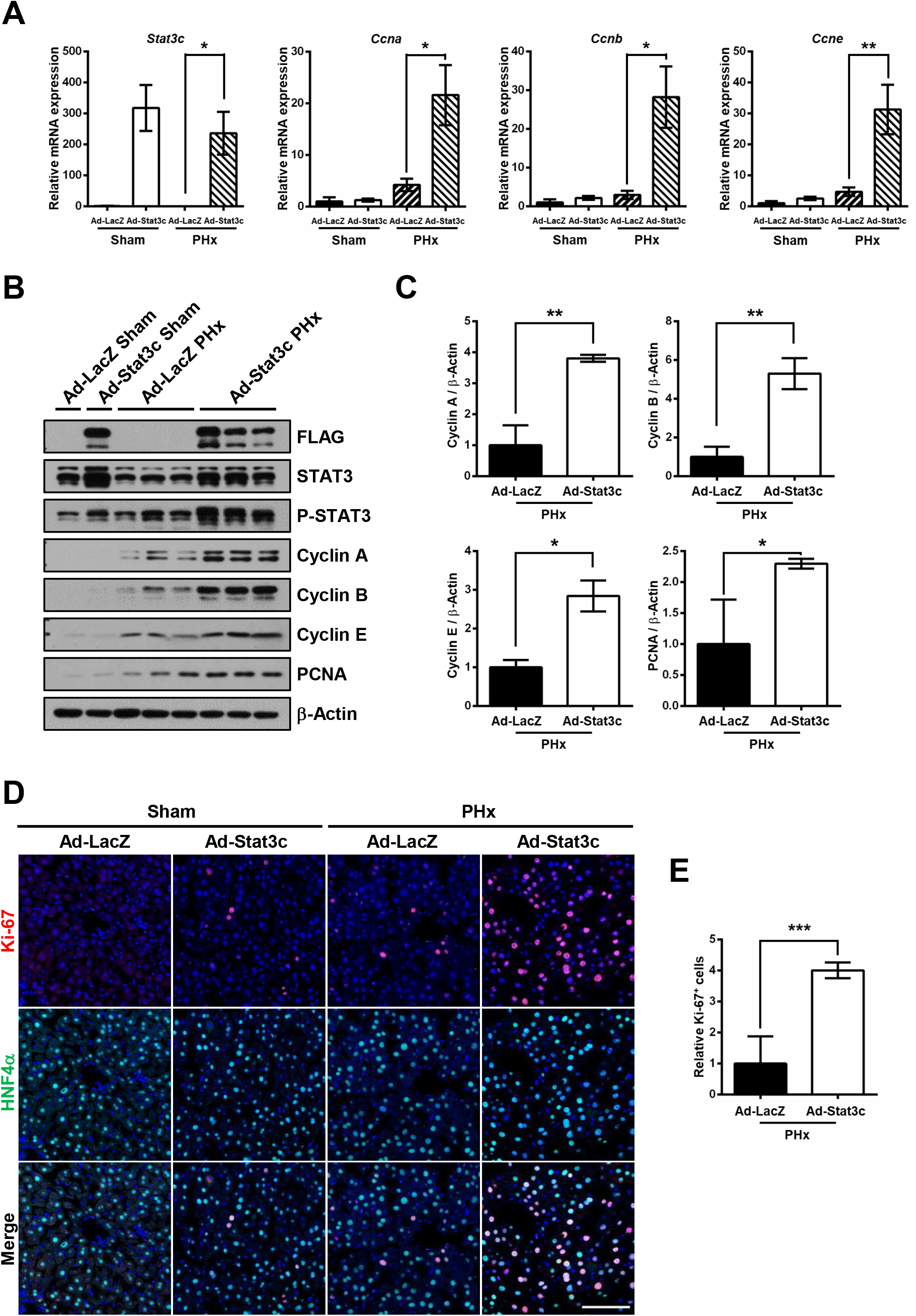
The expression of STAT3C rescues liver regeneration in liver-specific *Prom1*-deficient mice. A 2/3 partial hepatectomy was performed in 8-week-old male *Prom1^LKO^* mice after infection with adeno-LacZ (n=4-10) or adeno-Stat3c-FLAG (n=3-10). (A) The relative mRNA levels of *Stat3c*, *Ccne*, *Ccna*, and *Ccnb* in the liver 24 hours after PHx. Each mRNA level was normalized by 18S rRNA. (B, C) Immunoblotting for cyclin A, B and E, PCNA, FLAG, STAT3 and P-STAT3 in the liver 48 hours after PHx (B). Statistical analysis of the band intensities of cyclins A, B and E and PCNA in B. The band intensity of each protein was normalized to that of β-actin (C). (D) Double immunofluorescence for Ki-67 and HNF4α in the liver 48 hours after PHx. (E) Statistical analysis of the number of Ki-67-expressing cells (n=3). The number of Ki-67-positive cells was normalized to the number of DAPI-stained dots. Scale bar = 100 µm. Student *t*-test; \**p* < 0.05, \*\**p* < 0.01, \*\*\**p* < 0.001. Data are expressed as the mean ± SEM.

**Supplementary Table 1:**
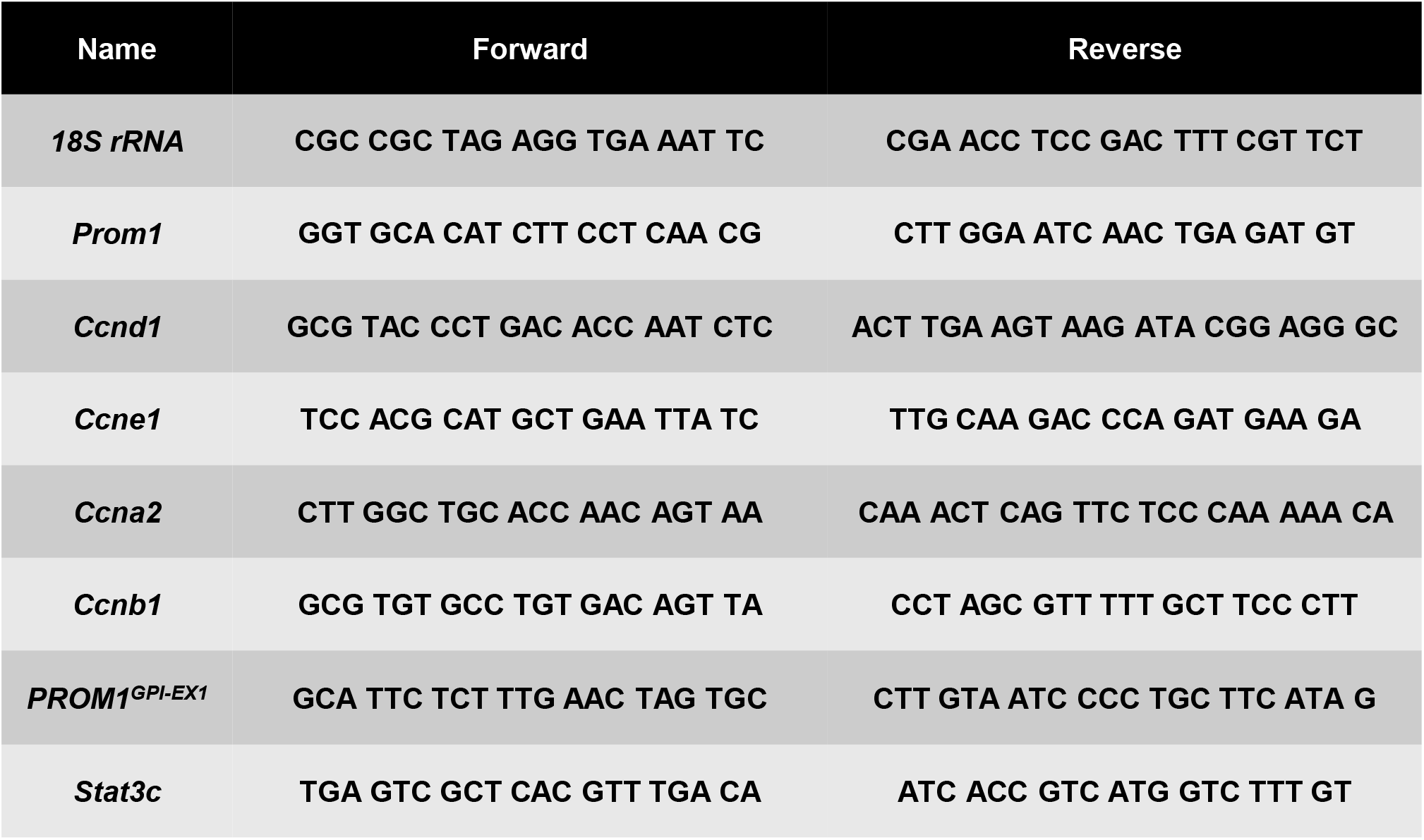

**Supplementary Table 2:**

| Name | Company | Host | Experiments |
| --- | --- | --- | --- |
| PROM1 | abcam | Rabbit polyclonal | IP |
| PROM1 | eBioscience | Rat monoclonal | IB (1:500) IF (1:100) |
| PROM1 | Developmental Studies Hybridoma Bank | Mouse monoclonal | IB (1:1000) |
| HNF4α | abcam | Mouse monoclonal | IF (1:200) |
| β-Actin | Santa Cruz Biotechnology | Mouse monoclonal | IB (1:1000) |
| CK19 | abcam | Rabbit monoclonal | IF (1:200) |
| Cyclin A | abcam | Rabbit monoclonal | IB (1:1000) |
| Cyclin B | Cell Signaling Technology | Rabbit polyclonal | IB (1:1000) |
| Cyclin E | Santa Cruz Biotechnology | Mouse monoclonal | IB (1:500) |
| Cyclin D | Santa Cruz Biotechnology | Mouse monoclonal | IB (1:1000) |
| PCNA | Cell Signaling Technology | Mouse monoclonal | IB (1:1000) |
| Ki-67 | Cell Signaling Technology | Rabbit monoclonal | IF (1:200) |
| P-STAT3 | Cell Signaling Technology | Rabbit monoclonal | IB (1:1000) |
| STAT3 | Cell Signaling Technology | Mouse monoclonal | IB (1:1000) |
| P-ERK | Cell Signaling Technology | Rabbit polyclonal | IB (1:1000) |
| ERK | Cell Signaling Technology | Rabbit polyclonal | IB (1:1000) |
| P-AKT | Cell Signaling Technology | Rabbit polyclonal | IB (1:500) |
| AKT | Santa Cruz Biotechnology | Rabbit polyclonal | IB (1:1000) |
| P-GSK3β | Cell Signaling Technology | Rabbit polyclonal | IB (1:500) |
| GSK3β | Cell Signaling Technology | Rabbit polyclonal | IB (1:1000) |
| FLAG | Merck Millipore | Rabbit polyclonal | IB(1:1000), IP |
| FLAG | Sigma-Aldrich | Mouse monoclonal | IB (1:2000) |
| GP130 | Cell Signaling Technology | Rabbit polyclonal | IB (1:1000) |
| Flotillin-1 | Santa Cruz Biotechnology | Rabbit polyclonal | IB (1:2000) |
| His | Santa Cruz Biotechnology | Mouse monoclonal | IB (1:1000), IP |
| RFP (tdTom) | Rockland | Rabbit polyclonal | IF (1:400) |
| tdTomato | Thermo | Mouse monoclonal | IF (1:400) |

## Notes

### Competing Interest Statement

The authors have declared no competing interest.

